# Attractor Landscapes as a Model Selection Criterion in Data Poor Environments

**DOI:** 10.1101/2021.11.09.466986

**Authors:** Cole A. Lyman, Spencer Richman, Matthew C. Morris, Hongbao Cao, Antony Scerri, Chris Cheadle, Gordon Broderick

## Abstract

Modeling of systems for which data is limited often leads to underdetermined model identification problems, where multiple candidate models are equally adherent to data. In such situations additional optimality criteria are useful in model selection apart from the conventional minimization of error and model complexity. This work presents the attractor landscape as a domain for novel model selection criteria, where the number and location of attractors impact desirability. A set of candidate models describing immune response dynamics to SARS-CoV infection is used as an example for model selection based on features of the attractor landscape. Using this selection criteria, the initial set of 18 models is ranked and reduced to 7 models that have a composite objective value with a p-value < 0.05. Additionally, the impact of pharmacologically induced remolding of the attractor landscape is presented.

## 1 Introduction

Even though broad-spectrum data is becoming increasingly accessible, biological modeling methods designed to operate in data poor environments are still critically important. The ongoing SARS-CoV-2 pandemic offers a highly relevant example of this where the research community has been called upon to make informed predictions of appropriate interventions when very little to no data is available, in particular molecular level data. Data-poor environments often result in underdetermined model identification problems, i.e. models for which there are more degrees of freedom than data-imposed constraints, which generally leads to multiple candidate models with equivalent adherence to a given objective. Under such conditions, multiple equivalently optimal sets of model parameter values exist. It is important however to consider the specific criteria used to define optimality of a given model. Adherence of model predictions to available data is a standard optimality criterion. Additionally, the complexity of a model is often minimized concurrently to achieve the most parsimonious adherence to data [36]. While adherence to data and parsimony are important criteria for assessing the desirability of a model, in cases where there are a large number of equally optimal models, i.e. equally data adherent and equally parsimonious, additional criteria may be required to further discern biological plausibility.

We propose that the number, type, and location of dynamically stable attractors (i.e. the attractor landscape) constitute useful criteria for further assessing the optimality of a regulatory network model. This is based on the widely accepted assumption that attractors can be used to represent phenotypes in models of biological systems [3,7,16,17,31,35]. In this work, we focus on qualitative models of immune regulation [20,44] and present methods to exhaustively compute attractors using this framework. Even though this work focuses on qualitative regulatory models, the ideas presented herein can be applied to any dynamical model type and are especially useful in mathematically underdetermined settings.

To demonstrate the utility of leveraging the attractor landscape to support additional optimality criteria, 19 distinct candidate models of the immune regulatory response to SARS-CoV [26] are analyzed. All competing candidate models reproduce the experimental data to within 5% departure and 11 of these match the available data exactly (100% accuracy). Furthermore, all of the models have a similar structural and parametric efficiency (i.e. structural and functional parsimony). The attractor landscapes, that is stationary steady states and limit cycles, for these 19 models are presented. These are compared in terms of relative abundance, location in state space, and oscillatory period along with their plausible biological counterparts. Moreover, the effects of potential repurposed drugs on these attractor landscapes are demonstrated suggesting that while some drugs serve to destabilize illness states, others also destabilize and make normal homeostatic regulation inaccessible.

### 1.1 Related Work

Identifying attractors in Boolean Networks has been shown to be an NP-complete problem in general [1, 50], but under certain conditions can be polynomially reduced to a satisfiability (SAT) problem [43]. Despite its NP-completeness, much work has been done to develop algorithms to efficiently characterize attractors. Some methods refine the state transition graph [2, 11, 21, 39], decompose the state transition graph into strongly connected components [25,48], or reduce the model itself [45], thereby limiting the search space and increasing efficiency. An alternative approach to efficiently identify attractors is to exploit SAT [10,14] and integer linear programming [32]methods. Moreover, some methods compute attractors by encoding the Boolean functions as binary decision diagrams [49, 51]. Finally, constraint satisfaction programming [9] has been utilized and is the paradigm upon which the current work is based.

Regardless of the multitude of methods proposed to identify attractors, to the best of the authors’ knowledge this work is the first to use the landscape defined by such attractors as a criterion for model selection.

## 2 Methods

This section briefly describes how 19 candidate regulatory models of immune response to SARS-CoV infection were obtained, and then proceeds with formal definitions of the model parameters and corresponding attractors the models support. Next, an algorithm for exhaustive attractor identification via constraint satisfaction programming is proposed. Finally, the criteria for evaluating a model’s performance is defined.

The network structure common to these 19 dynamic immune response models of SARS-CoV infection [26] consisted of 19 immune mediators (vertices) linked by 112 regulatory actions (edges) (see supplemental Fig. S1 for the network structure). These regulatory interactions were extracted from the Elsevier Knowledge Graph* (Elsevier, Amsterdam) [19] using the Pathway Studio* suite of tools [28] and were based on the automated text-mining [8, 29] of 2,653 published references. The system is modeled using a discrete dynamic network logic framework [20,44]. The predicted dynamic behaviors supported by any set of decisional logic parameters enacting these models were constrained to reproduce publicly available experimental measurements from time-series experiments (GEO accession number GSE33267) where human Calu-3 lung adenocarcinoma cells were infected *in vitro* by SARS-CoV over 72 hours with regular sampling for transcriptomic sequencing [38]. This dataset was chosen because high quality transcriptomic data of SARS-CoV-2 was not available at the time of analysis and SARS-CoV is closely related to SARS-CoV-2 [34]. The departure from these discretized data as well as structural and functional model complexity were concurrently minimized.

### 2.1 Attractor Definition

Let *A* be the set of attractors that a given model supports, i.e. the attractor landscape, where *a* ∈ *A* is an attractor. An attractor *a* is a set of states such that when a model reaches a state *a_i_* ∈ *a* the model will update to a state *a_j_* ∈ *a*. A state is a vector of length *m* where each element corresponds to a discrete value for an entity (vertex) in the model.

It follows that the number of states in *a* (|*a*|) is the period of the attractor, with the special case of |*a*| = 1 where the attractor is not a limit cycle (or oscillatory attractor) and can be referred to as a fixed stationary point, stable state, or steady state. An attractor can be viewed as a loop in the state transition graph where a steady state is a self-loop and a limit cycle of period *n* is a cycle of *n* states within the state transition graph.

### 2.2 Attractor Search Constraint Satisfaction Problem

Constraint Satisfaction Problems (CSP) are a class of optimization problems consisting of models that are composed of free variables, usually discrete, such that the permitted assignment of these variables satisfies all of the stipulated constraints that define the problem. A CSP solution is found when all variables are assigned and all the constraints are met. Here, the attractors for a model are formulated as a CSP and are encoded using the MiniZinc language (version 2.4.3) [27, 41] and the Chuffed solver (version 0.10.4) [6] to discover solutions (e.g. attractors). Pseudo-code of the CSP is found in the supplemental Algorithm S1.

The first variable defined in the CSP is *X*, a (*n* + 1) × *m* matrix that represents an attractor of period *n* where *n* > 0, *m* is the number of entities in the model, and where each row is a state within the attractor. The next variable is *I*, a (*n* + 1) × *m* matrix that represents the row-wise corresponding images for the states in *X*. The image *I* specifies the discrete values that the entities should move towards.

Next, constraints are stipulated to ensure that the states in *X* evolve according to the dynamics of the parameterized model. Formally, this constraint is 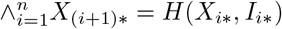 where *X_i*_* represents the *i*^th^ row of *X*, *I_i*_* represents the *i*^th^ row of *I*, and the *H* function computes the next state. The next state *X*_(*i*+1)*_ is determined by comparing the previous state *X_i*_* to the image *I_i*_* and updating entities according to the rules of the synchronous update scheme (see [5, 9, 36] for details on how the image computed and updates are performed in CSP). Also, the first and last states in *X* are constrained to be equal, thereby making *X* an attractor by definition. Formally, this constraint is *X*_1*_ = *X*_(*n*+1)*_.

An additional constraint only applies to cyclic attractors where *n* > 1, and constrains the first state in the cycle to not be equal to all of the other states excluding the last state. Formally, it is 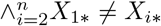 and ensures that the steady states (attractors where *n* =1) aren’t classified as limit cycles; because without this constraint, all steady states repeated *n* times would be valid solutions. The last constraint also only applies when *n* > 1 and ensures that the states in *X* are sorted in increasing order according to their sum. This constraint simply breaks the symmetry in the search space to prevent the same limit cycle from creating duplicate solutions. Without this constraint, each limit cycle would have *n* solutions where each solution has one of the *n* states as *X*_1*_. Formally, it is *X* = increasing(*X*) where the increasing function orders the rows of *X* increasingly by the L1-norm of the row.

Due to a technical limitation of MiniZinc which only allows static sized arrays, the variable *n* is treated as a parameter to the CSP. This means that separate searches for each *n* must be performed. The occurrence of attractors of period *n* tends to follow a power-law distribution [13] (also see Fig. 3b) with longer periods being increasingly rare. Thus, the search for attractors starts with *n* =1 and continues up to a pre-defined maximum period.

### 2.3 Pharmacologically Redirecting Attractors

Simulating how a drug will modify the attractor landscape is also accomplished as a CSP. First, the targets of a specific drug and the regulatory actions imparted to each of these targets are recovered based on data collected across multiple drug-action databases, including the Elsevier Reaxsys* database and integrated into the Elsevier Knowledge Graph*. In each case the targets are limited to those entities included in the model. Finally, an idealized drug action is applied such that it constrains the activation state of their corresponding targets in the network to a constant upregulated or downregulated state (assumption of continued administration) depending on that drug’s specific mode-of-action.

For a drug that is agonistic to a target (i.e. the drug up-regulates the target), the state value for that target is constrained to be at a level greater than its minimum level. For example, if a drug agonizes entity *v_i_* ∈ *V*, then *X_*i_* > 0 where *X_*i_* is the *i*^th^ column of *X*. Conversely, in the case of a drug that antagonizes a target (i.e. the drug down-regulates the target), the state value for that target is constrained to be less than its maximum level. This is formally represented as *X_*i_* < *ρ_i_* where *v_i_* ∈ *V* is the entity that the drug antagonizes and *ρ_i_* is the maximum level for *v_i_*. Note that this definition, when applied to a ternary target (*ρ_i_* = 2), allows for the state to have a nominal value of 1 indicating a smaller dose of the drug.

### 2.4 Model Performance Criteria

The formal definitions of the criteria used to select models are as follows. First, the adherence to data *O_d_* is calculated using (1) as the L1-norm ∥ · ∥_1_ (Manhattan distance) of the difference between the transient data *D* and the model-predicted trajectories *Tr* where |*D*| = |*Tr*| = *L* is the number of transient trajectories, *m* is the number of entities in the model, *Tr^i^* ∈ *Tr* is a *l_i_* × *m* matrix of the trajectory corresponding to the data *D^i^* ∈ *D*, and 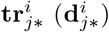 is the *j*^th^ row of *Tr^i^* (*D^i^*) and 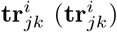 is the *k*^th^ element of the *j*^th^ row of *Tr^i^*(*D^i^*).

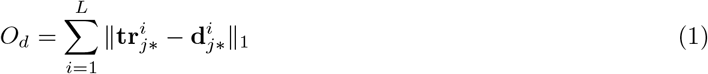

Optimal adherence to data is quantified as a minimal deviation of measured and predicted values, expressed as a fraction of the maximum possible error 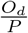, which accounts for absent data represented by ⊥ [36] and *ρ_k_* is the maximum level for *v_k_* ∈ *V*. The maximum level *ρ_k_* is used because it represents the largest possible amount of deviation between measured and predicted values when data is present.

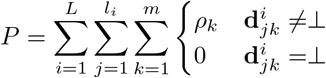

Additionally the model’s structural complexity *O_c_* should also be minimal and is computed here as the sum of the threshold values *ω* for all edges *E*, as illustrated in (2) (see [36] for further details on how complexity is calculated). Let (*i, j*) ∈ *E* be an edge from *v_i_* ∈ *V* to *V_j_* ∈ *V* with a corresponding activation threshold of *ω_ij_*.

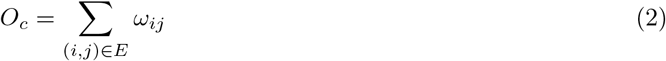

Turning now to the attractor space, the distance separating a given model’s attractors from a set of biologically relevant reference stable states *O_r_* should also be minimal. This is similarly calculated using a weighted sum of the L1-norm separating each reference state from the most proximal model-predicted attractors, as shown in (3). Let (*z_r_, r*) ∈ *R* be the set of reference stable states, e.g. stable health and persistent illness, where *r* is a reference stable state and *z_r_* is the associated weight of significance and let *a* ∈ *A* be an attractor in the set of attractors *A* for a given model. In this work the reference states are all equally weighted.

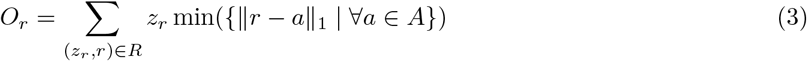

Third, we posit that the distribution of the periods corresponding to the cyclic attractors predicted by a given model should ideally favor higher frequencies, or in other words, be biased in favor of shorter periods. This translates to higher values of *O_p_*, as defined in (4) where |*a*| = *n* indicates that the attractor *a* ∈ *A* has a period of n, *N* = {|*a*| | *a* ∈ *A*} is the set of periods of the attractors in *A*, and *λ_n_*(*A*) is a function that returns the number of attractors in *A* with period *n*.

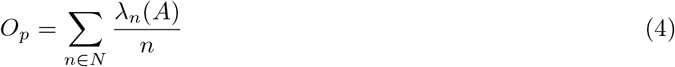

Finally the size of the basin, defined here (5) as the relative number of initial seed states that converge to that attractor, surrounding the non-pathological reference steady states *O_f_* should be maximal. Let *F*(*a*) be the relative frequency of convergence to attractor 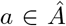 where 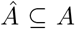 are attractors supported under non-pathological circumstances (e.g. no coronavirus present in the system) and where 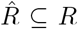 are non-pathological reference attractors.

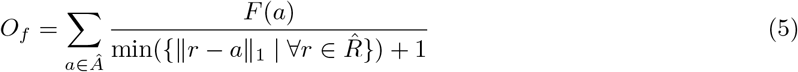

### 2.5 Model Selection

To rank the desirability of 19 models studied here based on their dynamic properties and the stable phenotypes they support, the 5 separate objective values in Section 2.4 are reduced to one composite value using Principal Component Analysis (PCA) [47]. The raw objective values for each objective are first individually scaled such that they range from 0 to 1. Additionally, objective values that should be maximal in the most suitable solution (*O_p_* and *O_f_*) are transformed so that the direction of each objective value agree. Error *O_d_* requires no scaling as it is already expressed as a percentage of the maximum error *P*. Structural complexity *O_c_* is scaled in terms of the maximum complexity value. Likewise, the adherence to reference stable states *O_r_* is scaled in terms of the maximum possible distance to the states. The reciprocal of the distribution of the periods *O_p_* is taken 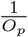, because *O_p_* value should be maximal. Lastly, the frequency of attractors most proximal to non-pathological reference steady states *O_f_* are filtered using add-one smoothing and the reciprocal is taken 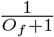. This transformation is performed because the minimum of *O_f_* is 0 and it should be maximized. Table 1 shows each objective value after their respective transformations.

**Table 1.** Objective Values

| Model ID | $O_d$ | $O_c$ | $O_r$ | $O_p$ | $O_f$ | PC1 Transformed Objective | P-Value |
| --- | --- | --- | --- | --- | --- | --- | --- |
| 18 | 0 | 0.7 | 0.019 | 0.19 | 0.83 | -2.5 | 2.5e-08* |
| 13 | 0 | 0.7 | 0.11 | 0.23 | 0.9 | -1.5 | 3.6e-05* |
| 17 | 0 | 0.7 | 0.019 | 0.2 | 0.96 | -1.4 | 1e-04* |
| 14 | 0 | 0.7 | 0.11 | 0.17 | 0.94 | -1.3 | 0.00017* |
| 19 | 0 | 0.7 | 0.11 | 0.12 | 0.97 | -1.2 | 0.00051* |
| 12 | 0 | 0.7 | 0.15 | 0.14 | 0.96 | -1.2 | 0.00055* |
| 15 | 0 | 0.7 | 0.019 | 0.26 | 1 | -0.83 | 0.0066* |
| 16 | 0 | 0.7 | 0.26 | 0.25 | 0.96 | -0.52 | 0.051 |
| 11 | 0 | 0.8 | 0.056 | 0.1 | 0.97 | -0.18 | 0.28 |
| 10 | 0 | 0.8 | 0.056 | 0.25 | 0.95 | 0.1 | 0.63 |
| 9 | 0 | 0.8 | 0.074 | 0.25 | 0.96 | 0.24 | 0.78 |
| 7 | 0.01 | 0.8 | 0.037 | 0.25 | 0.96 | 0.55 | 0.96 |
| 8 | 0.01 | 0.8 | 0.056 | 0.25 | 0.96 | 0.6 | 0.97 |
| 6 | 0.02 | 0.8 | 0.093 | 0.25 | 0.96 | 1.1 | 1 |
| 4 | 0.03 | 0.8 | 0.056 | 0.29 | 0.91 | 1.1 | 1 |
| 1 | 0.04 | 0.8 | 0.037 | 0.12 | 0.95 | 1.3 | 1 |
| 2 | 0.04 | 0.8 | 0.037 | 0.13 | 0.96 | 1.5 | 1 |
| 5 | 0.02 | 0.8 | 0.31 | 0.33 | 0.96 | 2 | 1 |
| 3 | 0.03 | 0.8 | 0.37 | 0.18 | 0.97 | 2.2 | 1 |

Following a PCA, the score values along the first latent feature PC1 (which explains 35% of the variation) are examined using the Shapiro-Wilk test to assess departure from normality [37] resulting in a p-value of 0.6935, suggesting PC1 scores describing an aggregate of the objective values do not depart significantly from a normal distribution (see Fig. 2a for a Quantile-Quantile plot of the distribution). Accordingly, a one-sided Student’s t-test [42] is performed to identify models having a significantly lower PC1 aggregate score. Table 1 lists the PC1 transformed objective values and the corresponding p-values for each model, where an asterisk indicates *α* ≤ 0.05. Furthermore, the PC1 transformed objective values with the corresponding Student’s t-test p-values are illustrated graphically in Fig. 2b.

## 3 Results

### 3.1 Model Selection

The original 19 SARS-CoV candidate models, all adhered to the experimental data to within 5% error with 11 of these matching the data exactly. A closer examination of the underlying regulatory complexity shows that 8 of these 11 models are less structurally complex (e.g. more parsimonious), where structural complexity is measured by the number and value of activation thresholds *ω* for each edge [36]. Conventional criteria of model error and parsimony would therefore leave us with 8 equally plausible models. We propose that these data-adherent parsimonious models could now be further ranked based on their attractor landscapes. We propose that such an extended model ranking scheme could not only favor 1. conventional adherence of model predictions to all available data, both steady state and transient, with minimal error and 2. structural and functional parsimony, but more importantly by also 3. biologically plausible predicted resting states that are maximally proximal to a set of stable *reference* states that are phenotypically relevant, 4. cyclic behaviors that favor shorter periods of oscillation, and finally 5. the accurate prediction of a reference non-pathological resting state with a broad basin of attraction as evidenced by high frequency of return after perturbation. As we require each model’s attractor landscape to demonstrate these features, we first conduct an exhaustive search of available stable states (*n* = 1) and limit cycles (1 < *n* ≤ 20) supported by each model. We find in this example that the limit cycles identified exhibit a maximum oscillatory period of *n* = 4. Details of the number of attractors with periods 1 ≤ *n* ≤ 4 found for each model are listed in Supplemental Table S1.

Now in addition to experimental data, we might also be able to define a number of dynamically stable *reference* states that correspond to a specific set of persistent biological phenotypes of interest, both pathological and non-pathological e.g. healthy homeostasis and chronic illness. Recovery of these *reference* states is represented by the *O_r_* objective. One could require that these *reference* states be recovered to within a user-specified tolerance as part of the attractor space supported by any feasible data-adherent model. Here we use 2 idealized phenotypes as reference states. First, we define a steady state of immune inactivation or immune quiescence to which the system should come to rest normally in the absence of an infectious or other challenge. This is idealized here as a state where all network entities exhibit a minimal level of activation, namely 0. In addition, we define a pathological state of persistent immune activation, namely cytokine storm, where network signaling mediators are expressed at high levels of activation. In this work, cytokine storm is modeled as the final state recorded in a 72-hour time course *in vitro* experiment, where viral titer had reached ~ 10^7^ pfu/ml and immune signaling activity was persistently elevated with respect to control. The Manhattan distance of the closest stationary steady state (*n* = 1) supported by each model to each of these reference steady states respectively is shown in Supplemental Table S1.

All of the 8 parsimonious, data-exact models support an attractor that is identical to the reference cytokine storm state. In contrast, the minimum distance to the reference state of normal immune quiescence is more varied, ranging from a Manhattan distance of 1 to 14.

Criterion (3), *O_r_*, was applied to these results, as a score consisting of a weighted sum of the minimal Manhattan distance to all reference steady states, with a lower score being more favorable. In this case the weights for each reference state are equal. We now have only 3 models where proximity to our set of reference stable attractors is equally well supported, namely models 18, 17, and 15. Under criterion (4), *O_p_*, we propose that the distribution of cyclic attractors is biased in favor of short oscillatory periods e.g. circadian or faster. This is expressed as a weighted sum of the number of attractors divided by their respective periods of oscillation *n*, with a higher score being more desirable. Results in Table 1 (and Supplemental Table S1) show that out of these 3 models, model 18 fares best in this regard with *O_p_* = 0.19, while model 15 does not support any oscillatory attractors and model 15 is biased towards supporting oscillatory behaviors of longer periods. This is illustrated further in Fig. 1 where the location of attractors for models 15 and 18 are presented. Each attractor exists in a 19-dimensional state space (19 state variables or network vertices in the model) that has been projected into 2-dimensions to facilitate visualization using Multidimensional Scaling [4] (implemented in Scikit-learn [15, 30]). In the case of oscillatory attractors (*n* > 1) arrows indicate the successor of each state.

**Fig. 1.**
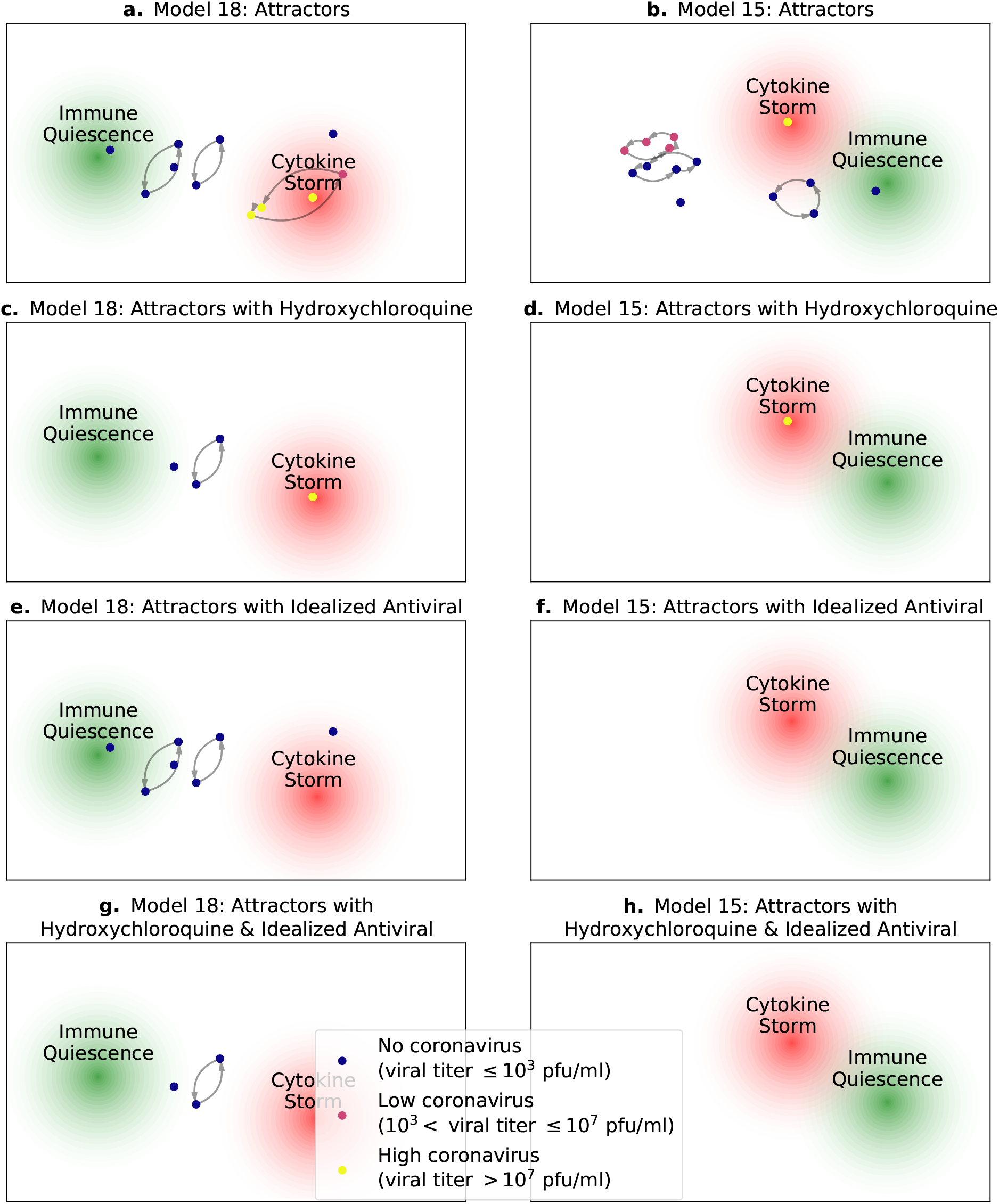
Attractors for models 18 and 15 with an idealized antiviral and hydroxychloroquine redirecting the attractors. **a, b.** The attractors for models 18 and 15 respectively with no drugs constraining the attractors. **c, d.** The attractors for models 18 and 15 respectively with hydroxychloroquine redirection, notice how the attractors near immune quiescence are no longer accessible for both models. **e, f.** The attractors for models 18 and 15 respectively with an idealized antiviral. **g, h.** The attractors for models 18 and 15 respectively with hydroxychloroquine combined with the idealized antiviral. Generated using Matplotlib [18].

Finally, under criterion (5), *O_f_*, we require that a non-pathological resting state (if defined) be the preferred steady state attractor for any system expected to support normal healthy homeostasis. Here we represent the breadth of the basin of attraction by the frequency with which the system returns to a given steady state in 100,000 random start states. The basins of attraction for the steady state attractors (*n* = 1) for models 15, 17, and 18 are shown in Table 2. These results indicate that in the absence of active infection, roughly 95% of the random perturbations to model 18 come to rest within a Manhattan distance of 6 bits or better from the reference resting state of immune quiescence. Indeed, 18% of the simulations conducted with model 18 come to rest in an attractor (attractor 2) located only 1 bit away from idealized immune inactivation. This is in stark contrast to models 15 and 17 where perturbations achieve such a final resting state proximity less than 1% of the time. This general proximity to a quiescent resting state is summarized in Table 2 by a composite score *O_f_* consisting of the sum across all attractors predicted in the absence of active infection (no coronavirus, i.e. viral titer ≤ 10^3^ pfu/ml) of their respective frequencies of occurrence divided by the corresponding Manhattan distances to the reference immune quiescent state. Finally, all 5 objective function scores are range-adjusted and combined into a single overall aggregate desirability using the first latent score from a Principal Component Analysis (PCA), with a lower aggregate score being preferable.

**Table 2.**
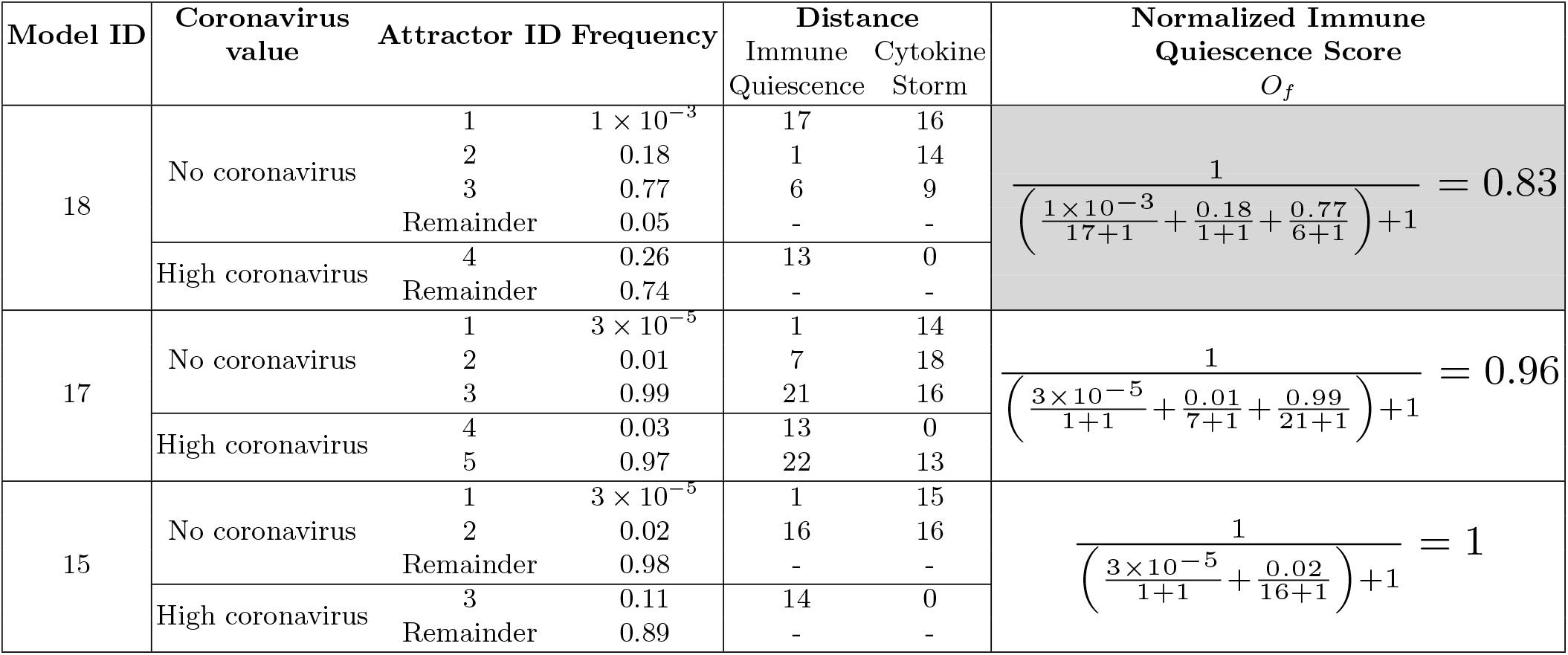
*n* =1 Attractor Frequency

### 3.2 Pharmacologically Redirected Attractors

Applying an idealized antiviral and the drug hydroxychloroquine to models 18 and 15, produces the attractor landscapes shown in Fig. 1c-h. This idealized antiviral is assumed to completely eliminate viral load, and its effects are approximated here by constraining the coronavirus activation level to its lowest value, 0. Fig. 2 panels f and h show that the simulated antiviral is sufficient to disrupt all attractors supported by model 15 and that hydroxychloroquine alone leaves intact only the virally fueled cytokine storm. In contrast, simulations conducted with model 18 tell a somewhat more plausible story. In Fig. 1c when only hydroxychloroquine is applied to model 18, it succeeds in disrupting the limit cycle that orbits cytokine storm as well as a normally occurring steady state proximal to it, leaving the virally supported cytokine storm stationary steady state in place. One quiescent limit cycle is also collapsed but a stationary steady state and another limit cycle proximal to the idealized immune quiescent state remain intact. In Fig. 1e the application of the idealized antiviral alone to model 18 leaves intact all limit cycles and stationary points that were proximal to “healthy” immune quiescence. However, it disrupts and renders unavailable the cytokine storm stationary point and limit cycle, leaving only one stationary point in the vicinity of cytokine storm. Lastly, in Fig. 1g we show the attractors supported by model 18 when both hydroxychloroquine and the idealized antiviral are applied. This combination regimen eliminates the cytokine storm steady state as well as all previously proximal pathological attractors, leaving the “parking” attractors proximal to immune quiescence as accessible stable resting states.

**Fig. 2.**
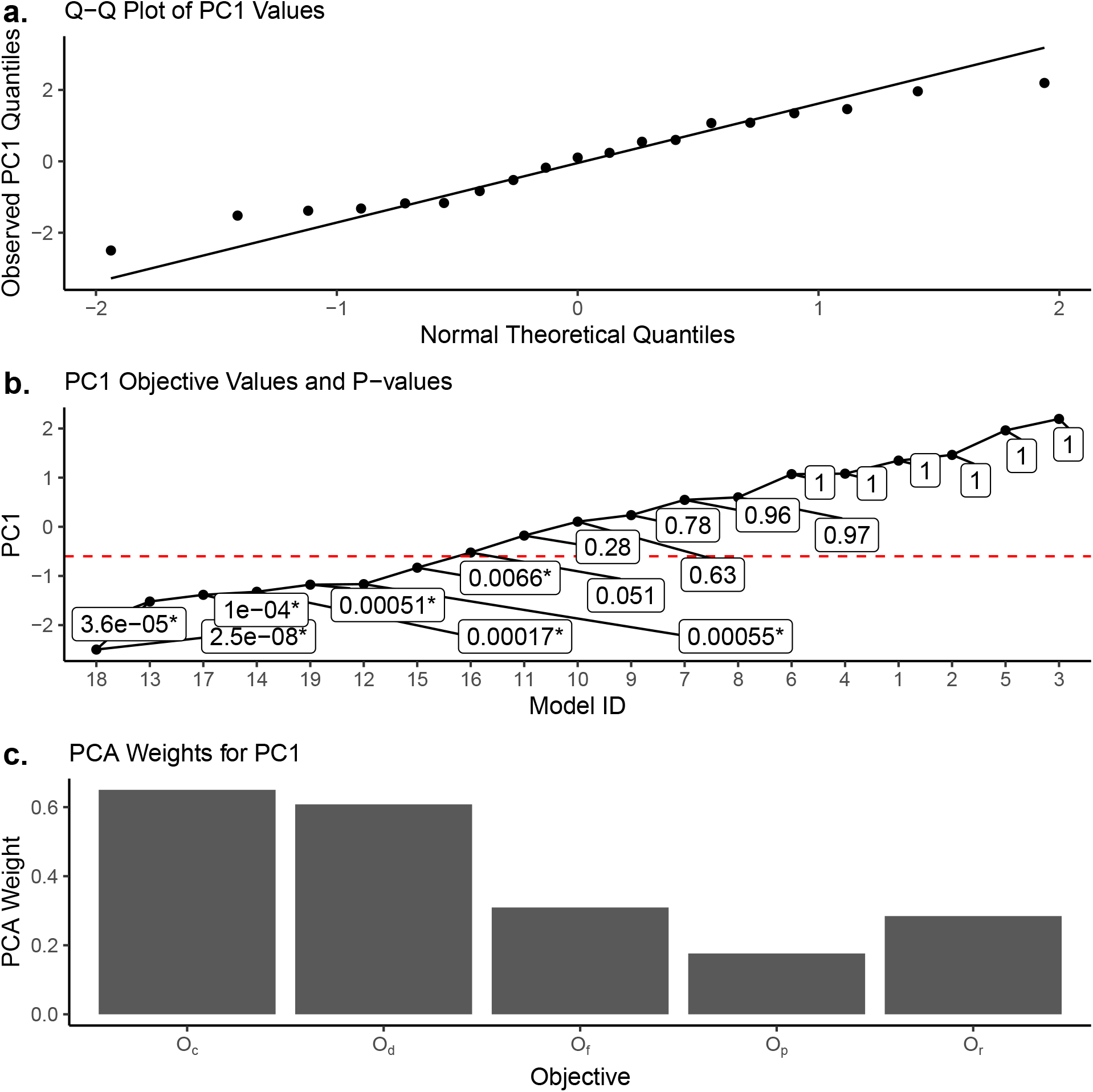
Dimensionally reduced objective values using Principal Component Analysis (PCA). **a.** A Quantile-Quantile (Q-Q) plot comparing the distribution of Principal Component 1 (PC1) objective values to the Gaussian distribution. The distribution of PC1 transformed objective values, with a Shapiro-Wilk [37] p-value of 0.6935 suggesting that these values do not depart significantly from a normal distribution. **b.** PC1 objective values for each model with associated p-values for a single sample, one sided Student’s t-test [42] where an asterisk indicates models with a significant p-value (< 0.05) and the dashed horizontal red line represents this significance threshold. **c.** The PCA weights for each objective. Generated using R [33], ggplot2 [46] and ggrepel [40].

### 3.3 Attractor Search Time Complexity

Searching for attractors is an NP-Complete problem [1, 50], thus the time complexity of the algorithm is an important consideration. Fig. 3a illustrates the mean time (in seconds) taken to search for attractors using model 18 in relation to the period of the attractor, n, where each search was repeated 10 times. Periods range from 1 to 20 (incrementing by 1) and from 20 to 90 (incrementing by 10) to provide a better sense of how the algorithm scales near-exponentially with regards to the period of the attractor. Despite the exponential scaling, searching for an attractor where *n* = 90 only takes approximately 8 minutes to determine that no such attractors exist. The whisker at each point is the standard deviation across the 10 repeats for that period. Fig. 3b illustrates the number of attractors found for that period, and shows that the attractors for model 18 follow a power-law distribution [13]. All experiments were run on a 2017 MacBook Pro with a 2.3 GHz Dual-Core Intel Core i5 processor and 8 GB of RAM.

**Fig. 3.**
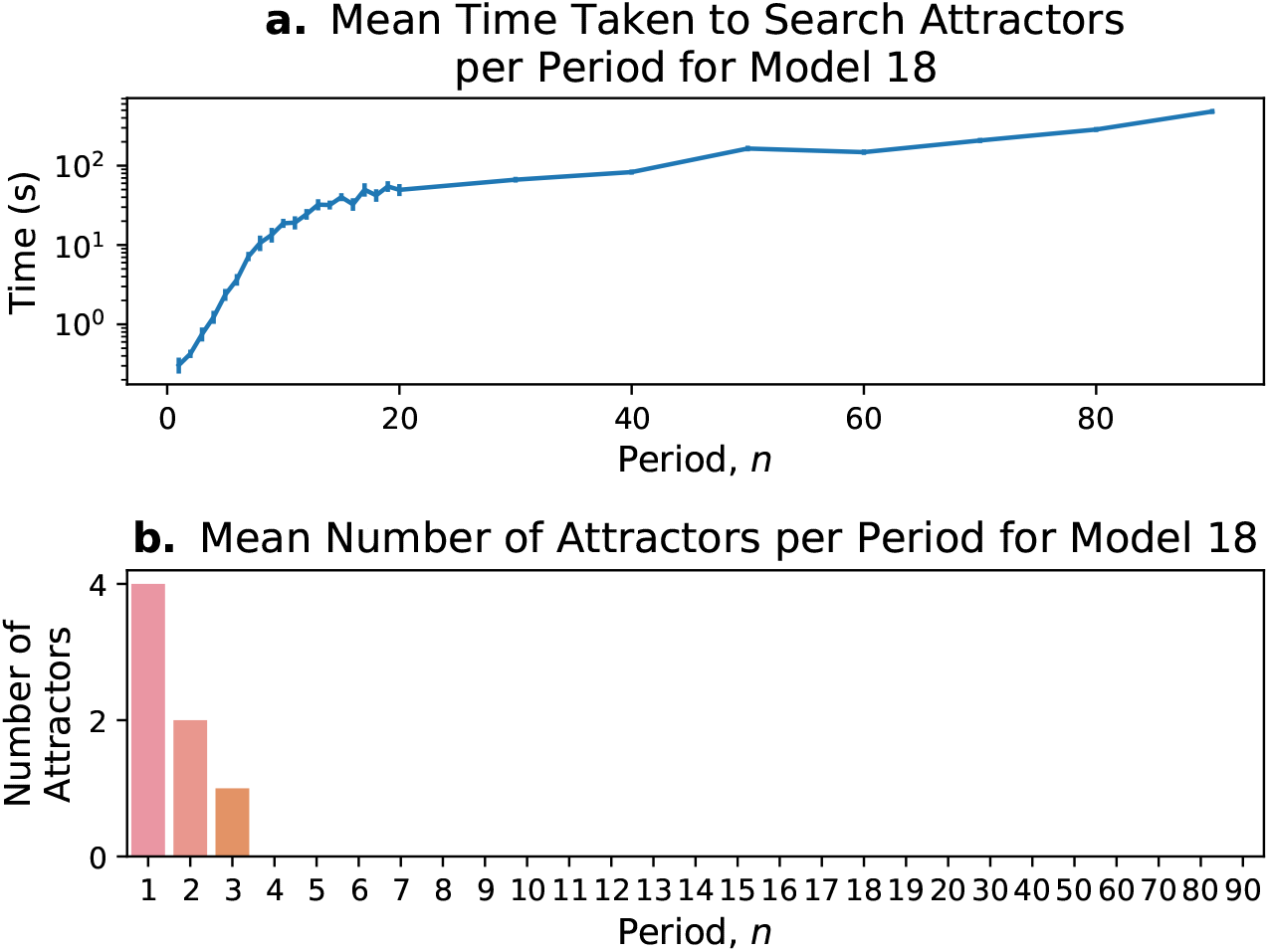
Time complexity and number of attractors per period. **a.** The mean time, in seconds, to search for attractors for model 18 repeated 10 times. Error bars represent the standard deviation for each period, which range from 1-20 and from 20-90 in increments of 10. **b.** The number of attractors identified for model 18 for each period, which follows a power-law distribution. Generated using Matplotlib [18].

## 4 Discussion

When selecting among competing models, we propose that in addition to adherence to reference data and model parsimony, the number, type, and location of attractors can offer important insight into biological plausibility. Moreover, we propose that the presence and periodicity of cyclic attractors is of special relevance to biological systems as these typically exhibit cyclic behaviors across multiple levels of resolution from the intracellular to entire organ systems [12]. Additionally, some illnesses have been shown to disrupt essential biorhythms which can be characterized as cyclic attractors, making such cycles of special therapeutic relevance.

It is interesting to note in the example presented here that hydroxychloroquine is predicted to not only disrupt the cyclic attractor proximal to cytokine store but also concurrently disrupt attractors nearest to immune quiescence. One might expect this behavior as strong immune mediators such as hydroxychloroquine aren’t meant to be administered for long periods of time, nor would they be administered as the sole agent in the case of cytokine storm [24]. Such results support ongoing efforts by our group directed at the development of computational methods for design of therapeutic approaches that formally account for the recovery of such essential cyclic attractors or biorhythms.

Given the time complexity exhibited by the current approach the authors are optimistic that other more complex cycles can also be characterized using this method. For example, this method could be applied and would be computationally feasible in the modeling of complex pulse-like oscillatory behavior to discover attractors such as those exhibited by the hypothalamic–pituitary–gonadal axis over the course of 30 days [23]. Furthermore, for less common biological phenomena that exhibit longer cycles, such as annual seasonal affective disorder [22], one could feasibly discover attractors with weekly measurements over the span of a year. In cases where an even larger number of states is needed to identify attractors efficiently the authors plan, as future work, to implement a flexible grid approach that allows for varying degrees of resolution in the attractor landscape.

## 5 Conclusion

This work presents a novel selection criterion of attractor landscapes that is useful in underdetermined modeling scenarios where models are equally optimal using conventional objective values. By combining conventional metrics, error and parsimony, with three attributes of the attractor landscape 18 models are ranked using a composite objective value. Out of the 18 original models, 7 models have a composite objective value that is significantly lower than the composite objective values of the other models. Using the top 2 ranking models, the effect of disrupting the attractor landscape with drug simulations is also shown. The attractor landscape is an important characteristic of network logic models and should thereby be included when selecting which models are suitable.

## Acknowledgment

This work was supported by Rochester Regional Health in conjunction with the US Department of Defense Congressionally Directed Medical Research Programs (CDMRP) http://cdmrp.army.mil/ under Peer Reviewed Medical Research Program (PRMRP) award W81XWH1910804 (Broderick - PI; Sethi - Partnering PI), as well as by Elsevier AB (Amsterdam). The authors would also like to thank Drs. Anton Yuriev, Philipp Anokhin, Dicle Hasdemir, Umesh Nandal, and Thibault Geoui of Elsevier BV for their technical assistance and many helpful discussions.

## Mandatory Disclaimer

The opinions and assertions contained herein are the private views of the authors and are not to be construed as official or as reflecting the views of the Department of Defense.

## Notes

### Competing Interest Statement

The authors have declared no competing interest.

