## Supplemental Figures for "Attractor Landscapes as a Model Selection Criterion in Data Poor Environments"

**Table S1.** Attractors Found for All Models

| **Model ID** | **Error** $\frac{O_d}{P}$ | **Structural Complexity** $O_c$ | **Minimum Distance** Immune Quiescence | Cytokine Storm | $O_r$ | **Number of Attractors** $n=1$ | $n=2$ | $n=3$ | $n=4$ | Total | $O_p$ |
|---|---|---|---|---|---|---|---|---|---|---|---|
| 18 | 0 | 0.7 | 1 | 0 | 1 | 4 | 2 | 1 | 0 | 7 | 5.3 |
| 17 | 0 | 0.7 | 1 | 0 | 1 | 5 | 0 | 0 | 0 | 5 | 5 |
| 15 | 0 | 0.7 | 1 | 0 | 1 | 3 | 0 | 1 | 2 | 6 | 3.8 |
| 19 | 0 | 0.7 | 6 | 0 | 6 | 6 | 4 | 1 | 0 | 11 | 8.3 |
| 14 | 0 | 0.7 | 6 | 0 | 6 | 5 | 2 | 0 | 0 | 7 | 6 |
| 13 | 0 | 0.7 | 6 | 0 | 6 | 3 | 2 | 1 | 0 | 6 | 4.3 |
| 12 | 0 | 0.7 | 8 | 0 | 8 | 6 | 2 | 1 | 0 | 9 | 7.3 |
| 16 | 0 | 0.7 | 14 | 0 | 14 | 3 | 2 | 0 | 0 | 5 | 4 |
| 11 | 0 | 0.8 | 3 | 0 | 3 | 6 | 6 | 2 | 0 | 14 | 9.7 |
| 10 | 0 | 0.8 | 3 | 0 | 3 | 4 | 0 | 0 | 0 | 4 | 4 |
| 9 | 0 | 0.8 | 4 | 0 | 4 | 4 | 0 | 0 | 0 | 4 | 4 |
| 7 | 0.01 | 0.8 | 2 | 0 | 2 | 4 | 0 | 0 | 0 | 4 | 4 |
| 8 | 0.01 | 0.8 | 3 | 0 | 3 | 4 | 0 | 0 | 0 | 4 | 4 |
| 6 | 0.02 | 0.8 | 5 | 0 | 5 | 3 | 2 | 0 | 0 | 5 | 4 |
| 5 | 0.02 | 0.8 | 17 | 0 | 17 | 3 | 0 | 0 | 0 | 3 | 3 |
| 4 | 0.03 | 0.8 | 3 | 0 | 3 | 3 | 1 | 0 | 0 | 4 | 3.5 |
| 3 | 0.03 | 0.8 | 16 | 4 | 20 | 4 | 3 | 0 | 0 | 7 | 5.5 |
| 1 | 0.04 | 0.8 | 2 | 0 | 2 | 6 | 5 | 0 | 0 | 11 | 8.5 |
| 2 | 0.04 | 0.8 | 2 | 0 | 2 | 6 | 3 | 0 | 0 | 9 | 7.5 |

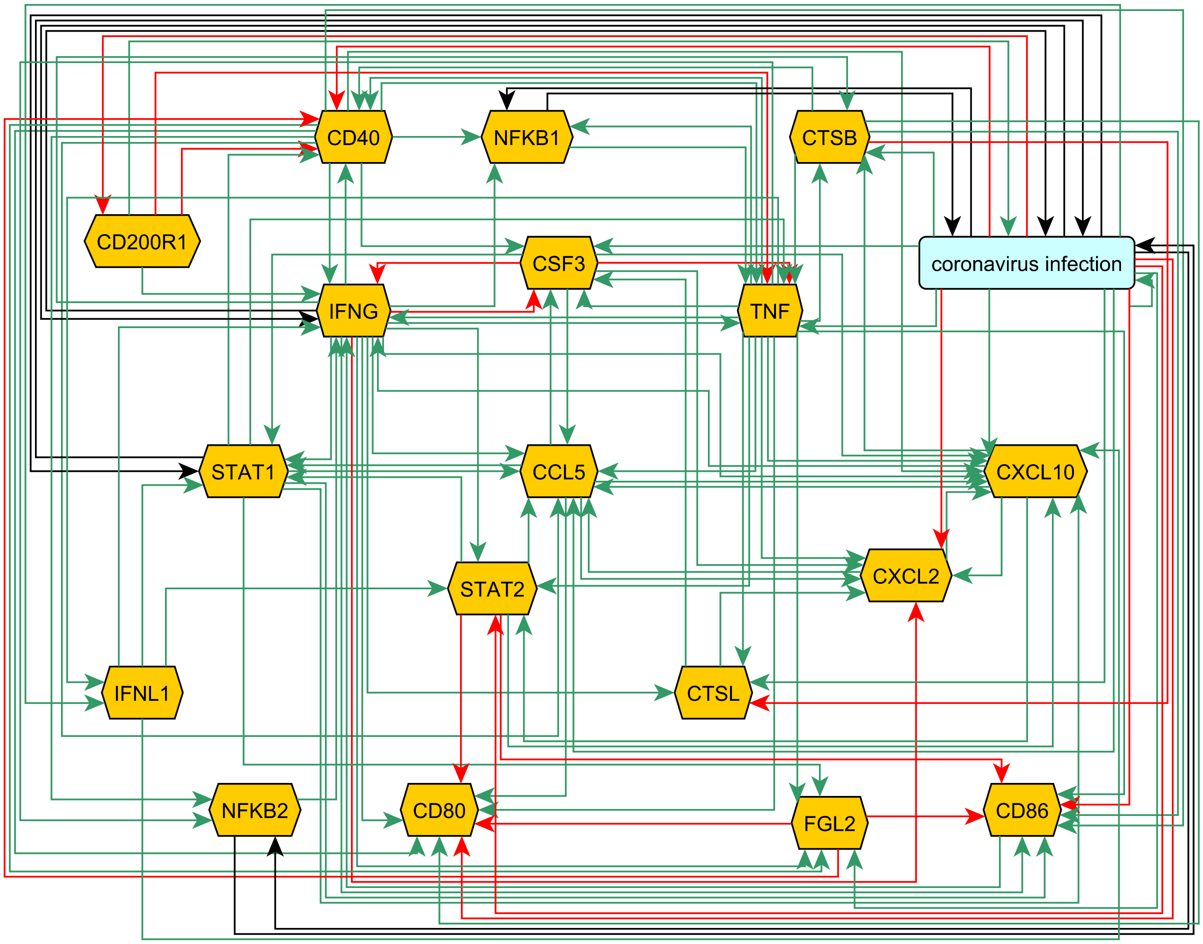


**Fig. S1.** Network structure of the SARS-CoV immune regulatory model [**?**]. The model contains 19 entites (nodes) and 112 regulatory actions (edges) which were obtained from biomedical literature using Pathway Studio*. The color of the edges indicates the mode-of-action for the regulation, where green is an activating mode-of-action, red is an inactivating mode-of-action, and black is an unknown mode-of-action. Generated using yEd [**?**].

*Copyright © 2021 Elsevier Limited except certain content provided by third parties. Pathway Studio is a trademark of Elsevier Limited.

**Algorithm S1** Attractor Search as a Constraint Satisfaction Problem

**Require:** $n$ is the period of the attractor
**Ensure:** $X$ is an attractor

**function** AttractorSearch($n$)
  $X \leftarrow (n+1) \times |V|$ matrix ▷ $|V|$ is the number of entities in the model
  $I \leftarrow (n+1) \times |V|$ matrix
  **for** $i \leftarrow 1$ to $n$ **do** ▷ combine the constraints in this block by logical conjunction (i.e. $\wedge$)
    $I[i,:] =$ ComputeImage($X[i,:]$)
    $X[i+1,:] = H^s(X[i,:])$ ▷ $H^s$ is the synchronous update function using $I$
    **if** $n > 1$ **then**
      $X[i+1,:] \neq X[i,:]$
    **end if**
  **end for**
  $X[1,:] = X[n+1,:]$ ▷ constrain the first and last row of $X$ to be equal
  $X =$ Increasing($X$) ▷ constrain the sum of each row in $X$ to be increasing
  **return** $X$
**end function**
